# Sampling Mismatch and Correction for Ptychographic Single-Particle Analysis

**DOI:** 10.64898/2026.02.21.707235

**Authors:** Tianyuan Li, Shouqing Li, Zhaoyi Yan, Yuan Shen, Xueming Li

## Abstract

Ptychographic single-particle analysis (SPA) is a promising technique for high-resolution biological imaging but is still limited by sub-nanometer resolution. In this study, we identified and investigated a critical issue termed sampling mismatch in ptychography that is caused by inaccuracies in the scanning step size and the pixel size of convergent beam electron diffraction (CBED) images. This mismatch induces pixel-size deviations in the reconstructed micrographs and modulates information transfer through a mismatch-induced modulation function (MIMF), which is characterized by phase reversals at specific spatial frequencies of the micrographs. These phase reversals, which vary with the defocus, cause destructive interference when merging micrographs, fundamentally limiting the resolution of SPA. We proposed a correction strategy and demonstrated, on the T. Acidophilum 20S proteasome and apoferritin datasets, that correcting sampling parameters eliminates signal distortions and improves resolution for ∼1.5 Å. These findings underscore the necessity for the precise control and calibration of the scanning system to achieve high-resolution ptychographic SPA.

## Introduction

Ptychography has recently emerged as a prominent technique to achieve sub-angstrom resolution^1^, even in uncorrected electron microscopes^2^, demonstrating significant potential for overcoming the resolution limitations of conventional methods such as phase-contrast imaging and scanning transmission electron microscopy (STEM). In this context, the integration of ptychography with cryo-electron microscopy (cryoEM) has attracted increasing attention^3–5^. However, the requirement for low-dose and large-area imaging of frozen-hydrated biological samples presents challenges for phase retrieval in ptychography. Consequently, although ptychography has been applied in single-particle analysis (SPA), commonly referred to as ptychographic SPA, its application has remained at a sub-nanometer resolution level, which is significantly lower than that achieved by conventional phase-contrast imaging^6,7^. Thus, elucidating the factors limiting ptychographic SPA has become imperative.

Prior to SPA, micrographs of frozen-hydrated biological specimens are generated through ptychographic phase retrieval from a series of convergent beam electron diffraction (CBED) images arranged on a STEM array. The quality of the micrographs, as well as the resolution of the subsequent SPA, critically depends on the information transfer from the electron optical system to the phase retrieval computation. Electron optical systems play a crucial role in extracting structural information and storing it in a series of CBED images. Phase-retrieval algorithms^8–11^, such as the extended ptychographic iterative engine (ePIE)^9^, process these CBED images along with a set of parameters that characterize the optical system, including the convergence semi-angle, probe defocus value, scanning step size, and CBED image pixel size (reciprocal-space pixel size). The accuracy of these parameters is essential for robust and high-quality phase retrieval. Although parameter deviations can, in principle, be corrected using optimization algorithms^12–15^, the practical performance of such algorithms is frequently limited by convergence to the local optima within a high-dimensional parameter space, which exhibits a strong dependence on the initial parameter values. This challenge may become more pronounced when the signal-to-noise ratio is low, particularly in low-dose biological specimen imaging, or in the presence of large aberrations in cryo-electron microscopes that typically operate without an aberration corrector. Furthermore, when combined with the requirements of biological samples for a large scanning step size and micrometre-level defocus, the estimation of initial parameters, such as the defocus and scanning step size, is often inaccurate. The influence of inaccurate parameters on ptychography for such biological applications has not been thoroughly investigated.

Here, we analyzed the deviations in the scanning step size and reciprocal-space pixel size of the CBED images, along with the defocus measurement in the presence of these deviations. We investigated their impact on information transfer in ptychographic SPA and proposed an effective correction strategy. A T. Acidophilum 20S proteasome dataset acquired using an uncorrected cryo-electron microscope and a published apoferritin dataset acquired using a corrected electron microscope^7^ were used to demonstrate the influence of inaccurate steps and pixel sizes on the resulting micrographs and final three-dimensional (3D) reconstructions. Note that the micrographs used here are imaginary parts of the transmission functions obtained by ptychography under a weak-phase-object approximation.

## Results

### Sampling mismatch in ptychography

The scanning step size (*δ*_s_) and the reciprocal-space pixel size (*δ*_f_) are two basic sampling parameters that indicate the sampling frequency in a ptychographic system. They are typically calibrated separately with independent deviations, which is problematic in ptychographic computations. To demonstrate the influence of these deviations, we first examined several explicit effects before proceeding to further analysis.

We consider a typical ptychography system with a detector of *N* × *N* pixels, a STEM array of *S* × *S* scanning positions, and a probe working at a defocus of *d*. To identify inaccurate parameters, we mark them with a tilde to denote values that may have possible deviations, referred to as nominal values. The nominal values for the two basic sampling parameters, the scanning step size 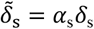 and the reciprocal-space pixel size 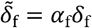, are described by the deviation factors, *α*_s_ and *α*_f_, from their true values, *δ*_s_ and *δ*_f_, respectively. Each CBED image recorded by the detector is processed in the ptychographic computation and contributes to the transmission function of the entire specimen in the real space as a patch image of *N* × *N* pixels with a nominal real-space pixel size of 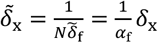, where 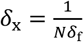 is the true real-space pixel size. The field of view of the transmission function has the nominal size of 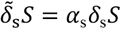 in each dimension defined by the STEM array. Using these nominal values as input, the ptychographic computation outputs a transmission function with 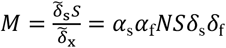 pixels in each dimension. Now, considering just the true size of the field of view, *δ*_s_*S*, and the number of pixels, *M*, output by the computation, the true real-space pixel size of the output is given as follows:

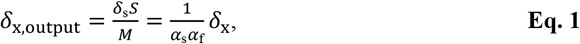

showing a modification to *δ*_x_ by a factor of 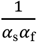.

The scanning step size (*δ*_s_) and pixel size (*δ*_f_ or *δ*_x_) are independently controlled in the microscope. The former is controlled by the scanning coils, and the latter corresponds uniquely to the camera length. Consequently, the deviation factors ( *α*_s_ and *α*_f_ ) should remain independent, yet ultimately combine to modify the pixel size *δ*_x_ by the factor 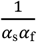, or simply, *α*_s_*α*_f_. To elucidate the factor *α*_s_*α*_f_, we examine the reconstruction step of the transmission function in various ptychographic algorithms, such as ePIE^9^, which integrates a series of partially overlapped image patches derived from corresponding CBED images (**Figure 1a**). In the ptychographic system, *α*_f_ affects the real-space pixel size of the image patches, whereas *α*_s_ affects the overlapping among these patches during computation. The deviations presented by *α*_f_ and *α*_s_ can collectively cause adjacent image patches to mismatch within their overlapping region, potentially producing ghosting artifacts (**Figure 1b**). However, ptychographic iterative optimization inherently compensates for such mismatches; therefore, the ghosting artifacts are not directly visible in the reconstructed micrographs (**Figure 1c**). The only observable effect is that the output pixel size *δ*_x,output_ deviates from its expected value 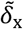. Briefly, the mismatch among the patches originates from the inconsistency of two basic sampling parameters, referred to as the sampling mismatch in this study, and leads to deviations in the pixel size, which ultimately affects the information transfer of the micrograph (discussed later). We defined *α*_s_*α*_f_ as the sampling-mismatch factor. We further analyze its impacts in subsequent sections.

**Figure 1.**
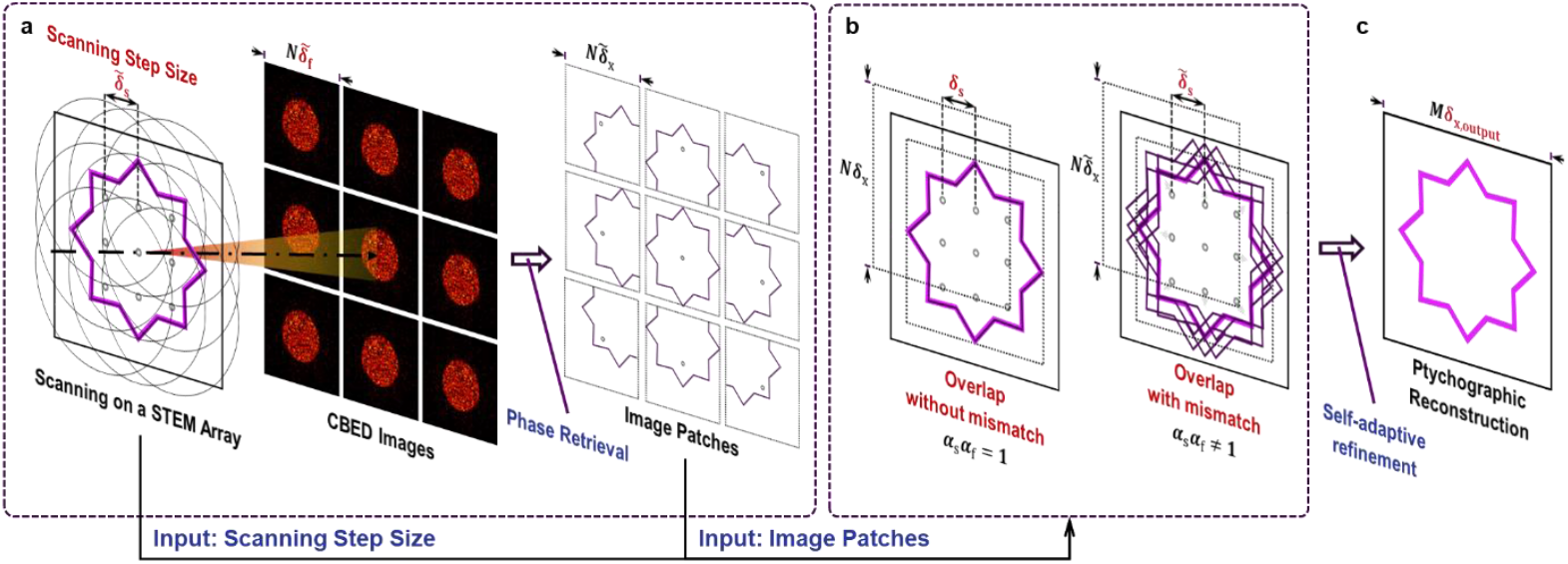
Schematic illustration of ptychographic reconstruction with and without sampling mismatch. **a**) The typical ptychographic processing, including scanning a sample (the magenta star) on a STEM array (the lattice of circles) and converting CBED images to image patches by phase retrieval. **b**) The ptychographic reconstruction illustrated by merging overlapped image patches with and without mismatch. **c)** The ghosting artifacts can be suppressed by the self-adaptive refinement.

The sampling mismatch can also affect the measurement of the probe defocus. The tilt-corrected bright-field STEM (tcBF-STEM) method^16^ is a powerful tool for estimating probe defocus in ptychographic workflows. In this method, a tcBF-STEM image can be constructed by combining the pixels at the same position *p* in all CBED images. These pixels correspond to the spatial-frequency *u* = *pδ*_f_ and the scattering angle *θ* = *λu* = *λpδ*_f_, where *λ* is the electron wavelength. The tcBF-STEM images have the pixel size *δ*_s_ and exhibit shifts among each other. The shift *s* = *tδ*_s_ for a tcBF-STEM image from the scattering angle *θ* is measured by *t* pixels relative to the tcBF-STEM image reconstructed from the CBED pixels at the zero scattering angle. Finally, the probe defocus is determined by 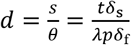. In the presence of deviations in the sampling parameters, the scattering angle of the CBED pixel has a nominal value 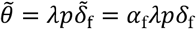, and the shift has a nominal value of 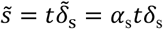. Replacing *s* and *θ* with their nominal values, respectively, yields the following estimated defocus

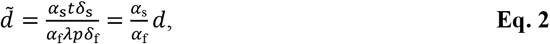

which shows that the measured value deviates from the true physical value *d* by a factor of 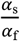.

In summary, the sampling mismatch impacts the ptychographic system in multiple aspects, as demonstrated by the factors, *α*_s_*α*_f_ and 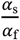.

### Size deviation and correction by single-particle analysis

An explicit effect induced by the sampling mismatch is the change in the output pixel size, which affects the nominal size of the protein particles in both the micrographs and reconstructed 3D density maps. We examined this effect with two datasets in ptychographic SPA.

The T20S proteasome dataset was acquired with the nominal sampling parameters, a scanning step size of 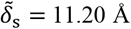 in a 300 × 300 STEM array, and a reciprocal-space pixel size of 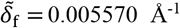 (corresponding to 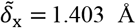 in real space). Ptychographic micrograph reconstructions were performed using a batched stochastic gradient descent algorithm^17^ implemented in a custom MATLAB pipeline (See **Methods** and **Code Availability**). We then selected particles from those micrographs with a size of 2387 × 2387 pixels (**Supplementary Figure 1a**) and subjected them to 3D reconstruction using THUNDER^18^ (**Figure 2a**). Subsequently, we observed that the reconstructed density map exhibited poor fitting with the atomic model (PDB entry code: 1PMA^19^) in terms of size, indicating the existence of a pixel-size deviation, as previously discussed. To correct the size problem, we fitted the reconstructed map to the PDB model using the EMDA software package^20^ (See **Methods**), and calibrated the pixel size as *δ*_x,output_ = 1.278 Å. After updating the pixel size, the reconstructed map exhibited a shrinkage of 9.8%, which matched well with the atomic model (**Figure 2a**). Accordingly, we obtained the deviation factor 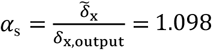, subsequently, we achieved a corrected scanning step size *δ*_s_ =10.20 Å. Using the corrected scanning step size together with a pixel size of 1.403 Å, the size of the micrographs was updated to 2175 × 2175 pixels by recalculating the ptychographic reconstructions (**Supplementary Figure 1b**). Note that the pixel size of 1.403 Å is thought to be the true value (*α*_f_ = 1.00), which has been calibrated using the electron diffraction of a diffraction standard sample of Evaporated Aluminum (TED PELLA, INC., Product No. 619).

**Figure 2.**
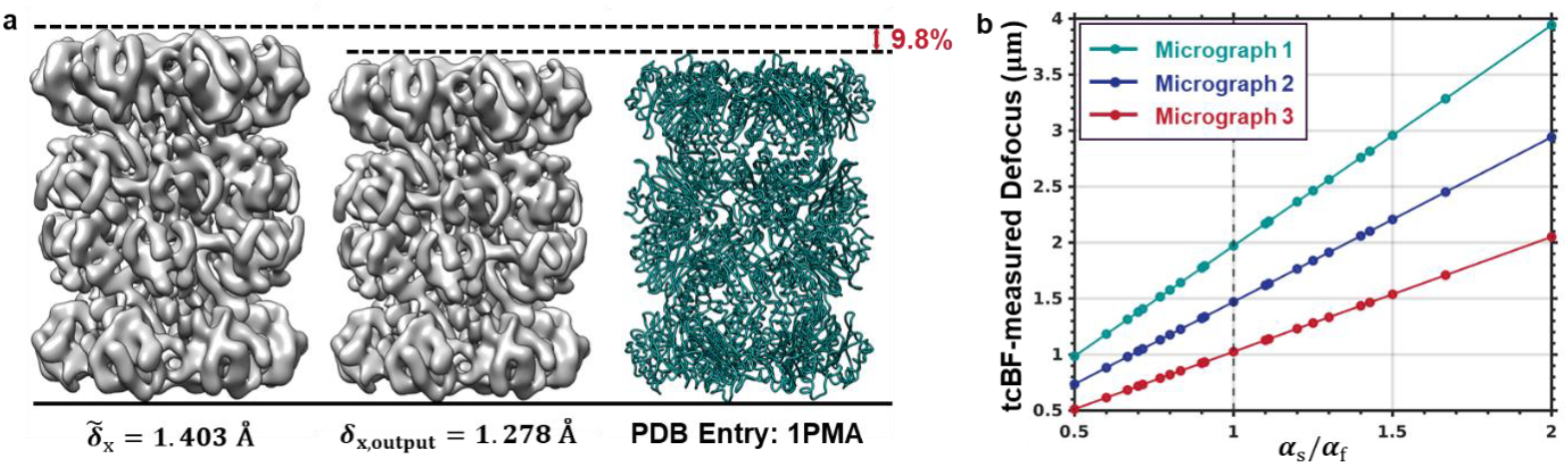
Output size and defocus measurement influenced by the sampling mismatch. **a)** Comparisons of the reconstructed 3D density maps shown using the nominal (1.403 Å) and corrected (1.278 Å) pixel size, with the atomic model. **b**) The distribution of the defocus measured by the tcBF-STEM method against *α*_s_/*α*_f_ . Each dot represents an independent measurement, and the measured defocuses of three micrographs respectively distributed on three lines.

We measured the defocus using the tcBF-STEM method and obtained different values for various deviations in the sampling parameters. As expected, the ratio between the defocuses measured using the nominal parameters and the newly calibrated scanning step size was *α*_s_/*α*_f_ = 1.098. Furthermore, through additional testing by varying *α*_s_ to simulate different *α*_s_/*α*_f_ ratios, we observed that this ratio remained strictly consistent with *α*_s_/*α*_f_ (**Figure 2b, Supplementary Figure 2**), thereby confirming the conclusion presented in the previous section.

The same tests (data not shown) as above were also applied to the apoferritin dataset^7^, which gave the factor *α*_s_ = 0.971, indicating a deviation of 2.9% in the scanning step size from 20.00 Å to 20.60 Å in a 128 × 128 STEM array. This further indicates that deviations in scanning step size are common.

The influence of the sampling mismatch was not limited to misinterpretation of the output pixel size and measured defocus. This may also cause information modulation in the ptychographic algorithms. After correcting for the scanning step size, we observed resolution improvements in the ptychographic SPA, which are discussed in detail in the last section.

### Phase reversal induced by sampling mismatch

To observe the possible information modulation induced by the sampling mismatch, we compared the signal differences between micrographs affected and unaffected by the sampling mismatch. A comparison was performed by calculating the Fourier ring correlation (FRC), defined as

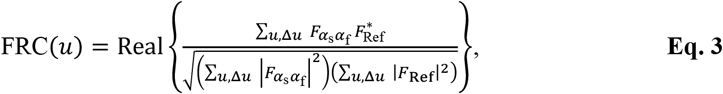

where *u* denotes the spatial frequency, ∑_*u*,Δ*u*_(·) indicates summation over all points within the ring region at spatial frequency *u* with radial thickness Δ*u*, the symbol ∗ denotes the complex conjugate, and Real{·} indicates the real part. *F*_*α*s_*α*_f_ and *F*_Ref_ are the Fourier transforms of micrographs affected (with deviation factors *α*_s_ and *α*_f_ ) and unaffected (reference) by the sampling mismatch, respectively.

For comparison, the reciprocal-space pixel size and scanning step size of the T20S proteasome and apoferritin datasets were calibrated based on 3D reconstruction, as described previously, and considered as the true values. Accordingly, the micrographs calculated using the true sampling parameters were used as references. Subsequently, a series of micrographs were generated with different combinations of *α*_s_ and *α*_f_ and subjected to the comparison with the corresponding references. Because the sampling mismatch makes the pixel size and the object size deviate from their true values, to enable a comparison with the reference micrographs, the true pixel sizes, *δ*_x,output_, were calculated and used to calculate the FRC for the micrographs affected by the sampling mismatch. As shown in **Figure 3**, remarkable effects induced by the sampling mismatch were observed in the comparison.

**Figure 3.**
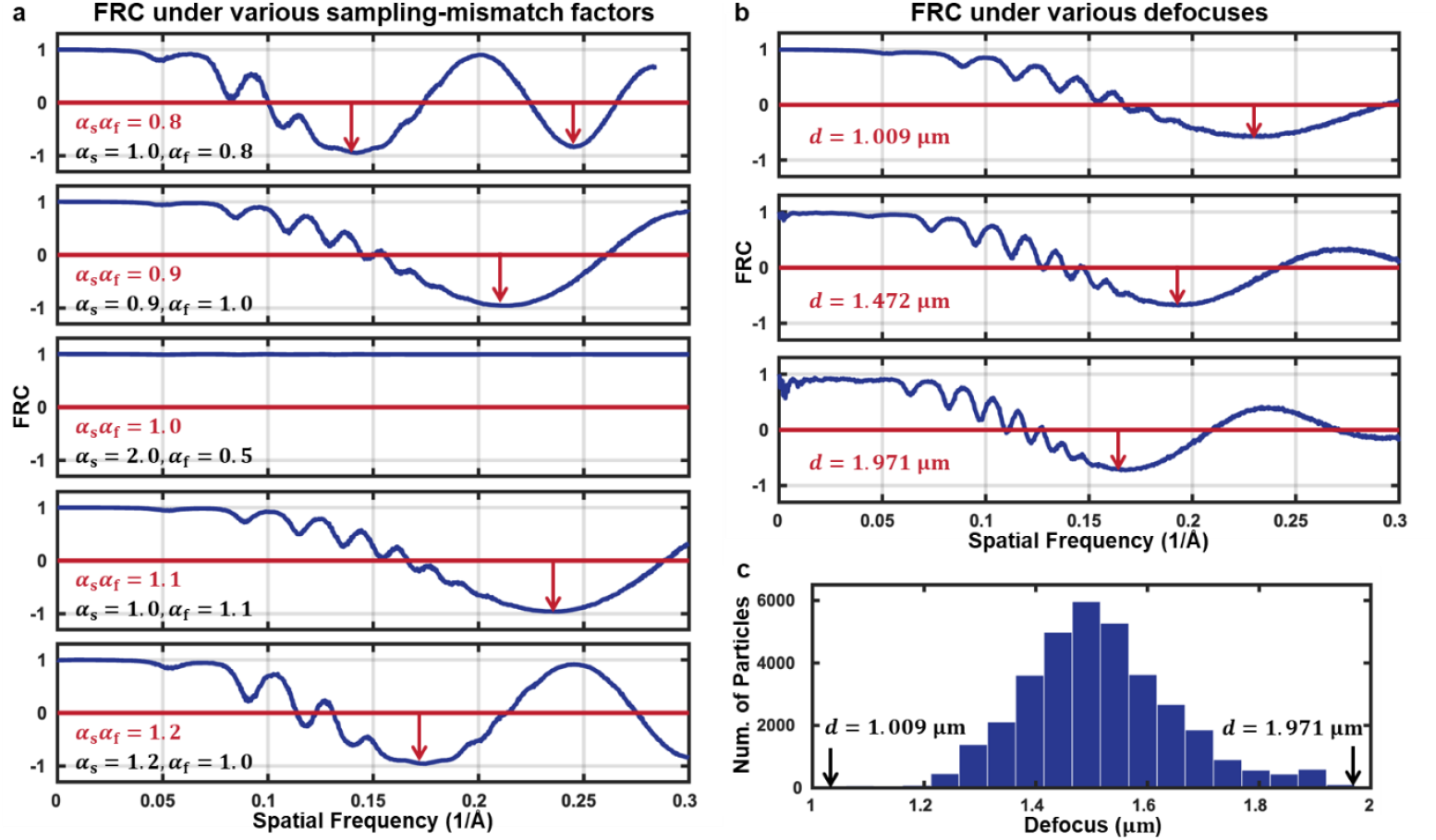
FRCs under various sampling-mismatch factors and defocus. **a**) FRCs under various sampling-mismatch factors (with the same true physical defocus). **b**) FRCs under various defocuses (with the same sampling-mismatch factor). **c**) Histogram of the defocus values of all particles in the T20S proteasome dataset.

First, the FRCs change with *α*_s_*α*_f_ and exhibit an overall cosine-like oscillation combined with fast fluctuations at low spatial frequency (**Figure 3a**). Intriguingly, FRCs remain unchanged for any combinations of *α*_s_ and *α*_f_ if *α*_s_*α*_f_ is the same (the defocus should be accordingly amended according to **Eq. 2**) (**Supplementary Figure 3**). This means that the deviations of the CBED-image pixel size and the scanning step size act collectively on the ptychographic reconstruction through the product of *α*_s_*α*_f_ . Furthermore, the value of *α*_s_*α*_f_ presents the level of the mismatch. When *α*_s_*α*_f_ = 1, the FRC is always 1, indicating that the ptychography give an output that was the same as that with accurate sampling parameters (**Supplementary Figure 3b**), implying the two independent experimental systems, the acquisition for the CBED images and the scanning control, match each other even though *α*_s_ and *α*_f_ are not 1. With increasing deviation of *α*_s_*α*_f_ from 1, the cosine-like oscillation increases continuously (**Figure 3a**), indicating the increasing mismatch.

Second, these cosine-like oscillations introduce zero crossings, indicating the presence of phase reversals in the spatial frequency where FRC < 0 (red arrows in **Figure 3a and b**). By checking the FRCs calculated from the micrographs with different defocuses in the T20S proteasome dataset, we observed that the phase reversals varied with defocus (**Figure 3b**). While the value of *α*_s_*α*_f_ might be unchanged in a session of data acquisition, the defocus is usually inclined to vary (**Figure 3c**). Similar to the influence of the contrast transfer function (CTF) in traditional SPA, phase reversals lead to signal attenuation when images with different phase-reversal patterns are combined. This attenuation ultimately limits the resolution of the resulting 3D reconstructions.

In summary, sampling mismatch (*α*_s_*α*_f_ ≠ 1) induces phase reversals that vary with *α*_s_*α*_f_ and the defocus, thereby imposing a fundamental resolution limit in ptychographic SPA.

### A Fresnel-propagation model of phase reversal

To formulate the phase reversal, we hypothesized the following Fresnel-propagation model of the sampling mismatch. According to the discussion in the previous section, the phase reversals induced by sampling mismatch depend solely on the product of *α*_s_*α*_f_ and the true defocus *d*. Consider a CBED image dataset acquired with true defocus *d*, processing it with a deviation factor set of (*α*_s_, *α*_f_) and the corresponding nominal defocus 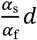 is equivalent to processing it with a set of (*α*_s, eq_ = 1, *α*_f, eq_ = *α*_s_ *α*_f_ ) and the corresponding nominal defocus 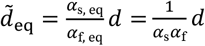. Here, we set *α*_s, eq_ = 1 to attribute all effects of the sampling mismatch to the optical diffraction system with the pixel-size deviation of CBED images. Accordingly, the sampling mismatch changes the defocus from *d* to 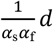. In an optical system, a defocus change usually implies an extra propagation of the wave function. Therefore, if regarding the ptychographic reconstruction as a virtual optical system, the calculated transmission function 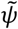 might be interpreted as the true transmission function *ψ* after undergoing free-space propagation for an additional distance of 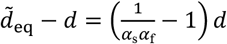 (**Figure 4a**). Subsequently, we have

**Figure 4.**
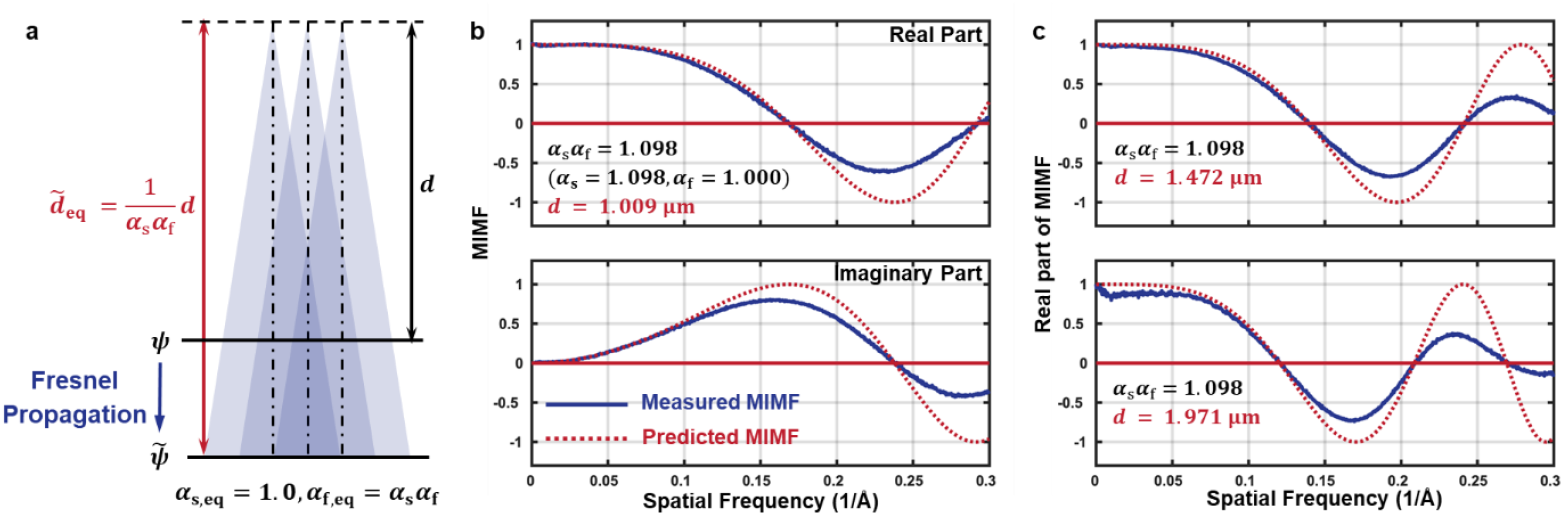
Model and validation of the mismatch-induced information modulation function. **a**) Schematic illustration of the defocus change and Fresnel propagation induced by the sampling mismatch. **b**) The measured MIMF under *α*_s_*α*_f_ = 1.098 and the true defocus of 1.009 μm. **c**) The real part of the measured MIMF under true defocuses of 1.472 μm and 1.971 μm for the same *α*_s_*α*_f_ = 1.098. The solid blue lines and the red dashed lines indicated the measured and predicted curves, respectively.

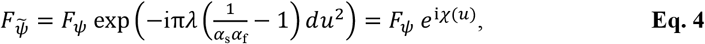

where 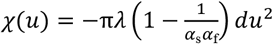 and the exponential term presents a Fresnel propagation. *F*_*ψ*_ and 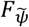 denote the 2D Fourier transforms of *ψ* and 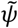, respectively. Here, we define *e*^i*χ*(*u*)^ as the mismatch-induced modulation function (MIMF) that describes information modulation in the presence of the sampling mismatch.

To validate the MIMF described in **Eq. 4**, we processed a set of CBED images in the T20S proteasome dataset with and without sampling mismatch, respectively, yielding a pair of transmission functions, 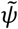 (affected) and *ψ* (unaffected). The information modulation between *ψ* and 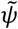 was then estimated using a least-squares formulation (See **Methods**). As shown in **Figure 4b**, the estimated modulation function agrees well with the hypothesized MIMF, in addition to some differences in the amplitude attenuation. Further comparisons across different sampling-mismatch factors *α*_s_*α*_f_ (**Supplementary Figure 4**) and defocus values (**Figure 4c**) also confirmed the MIMF.

Furthermore, a FRC formula can be derived from MIMF, as

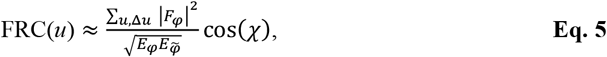

where *φ* and 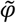 are the projection potential functions that are unaffected and affected by the sampling mismatch, respectively, and 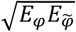 presents the noise level in the Fourier ring at spatial frequency *u* under the low-dose condition (see **Methods**). The cos(*χ*) captures the phase-reversal behavior induced by sampling mismatch and explains the overall oscillation in the FRC curves (**Figure 3**). The fluctuations superimposed in the low frequency region of FRC (**Figure 3**) should be from the intrinsic properties of the underlying signal of *φ*.

### Improved resolution in ptychographic single-particle analysis

Size deviations and phase reversals have been identified as being related to sampling mismatch that is mostly caused by inaccurate sampling parameters. Although the size deviation itself is a concern, phase reversals represent a more critical issue in SPA. In SPA, particle images are typically acquired with varying defocuses. In the presence of a sampling mismatch, these varying defocus values lead to distinct phase-reversal patterns among the particles. Averaging such particles introduces signal degradation. To evaluate the impact of phase reversals quantitatively, we assessed both the attainable resolution and map quality using the T20S proteasome and apoferritin datasets.

For each dataset, two sets of micrographs were generated based on the uncorrected and corrected sampling parameters as described in the previous section. The defocus value of each micrograph was determined using tcBF-STEM with the corresponding sampling parameters. Subsequently, each set of micrographs was processed independently using SPA. Notably, the corrected true pixel size *δ*_x,output_ was applied to the micrographs during SPA to ensure that the resulting density map maintained correct size. Ultimately, ∼100 k particles from 388 micrographs of the T20S proteasome dataset and ∼22 k particles from 73 micrographs of the apoferritin dataset were selected through 2D classifications and subsequently used for 3D reconstructions with THUNDER^18^ (See **Methods**).

The reconstructions demonstrated marked improvements between the two sets of micrographs (**Figure 5 and Table 1**), hereafter referred to as the corrected and uncorrected reconstructions. The Fourier shell correlation (FSC) curves revealed resolution enhancements exceeding 1.5 Å (**Figure 5a**). Notably, abnormal oscillations yielding negative FSC values in the 4-6 Å resolution range were observed in the uncorrected reconstructions for both datasets, whereas these artifacts were absent in the corrected reconstructions. Similar FSC oscillations were detected during processing using both CryoSPARC^21^ and RELION^22^ (**Supplementary Figure 5**), indicating that these oscillations originated from anomalous signal characteristics rather than from artifacts induced by the processing software. The resolution range showing abnormal oscillations approximately corresponds to the spatial-frequency range of the first zero crossing of the MIMF (gray band in **Figure 5a**), suggesting a direct correlation with the sampling mismatch.

**Table 1.** Map statistics from different software packages and datasets with corrected and uncorrected sampling parameters.

| Dataset Name<br>Softwares |  | Apoferritin |  | T20S Proteasome |  |
| --- | --- | --- | --- | --- | --- |
|  |  | Uncorrected | Corrected | Uncorrected | Corrected |
| **THUNDER** | Corrected Pixel Size (Å) | 1.545 | 1.500 | 1.278 | 1.403 |
|  | Number of Particles | 22,732 | 25,665 | 102,966 | 105,665 |
|  | B-factor (Å<sup>2</sup>) | -291 | -155 | -380 | -227 |
|  | Resolution (Å) | 7.51 | 5.85 | 7.79 | 6.30 |
| **CryoSPARC** | Corrected Pixel Size (Å) | 1.545 | 1.500 | 1.278 | 1.403 |
|  | Number of Particles | 20,853 | 24,687 | 97,443 | 104,884 |
|  | B-factor (Å<sup>2</sup>) | -169 | -546 | -201 | -129 |
|  | Resolution (Å) | 7.03 | 5.46 | 7.98 | 5.89 |
| **RELION** | Corrected Pixel Size (Å) | 1.545 | 1.500 | 1.278 | 1.403 |
|  | Number of Particles | 22,732 | 25,665 | 102,966 | 105,665 |
|  | B-factor (Å<sup>2</sup>) | -221 | -628 | -237 | -182 |
|  | Resolution (Å) | 7.27 | 6.00 | 7.61 | 6.78 |

**Figure 5.**
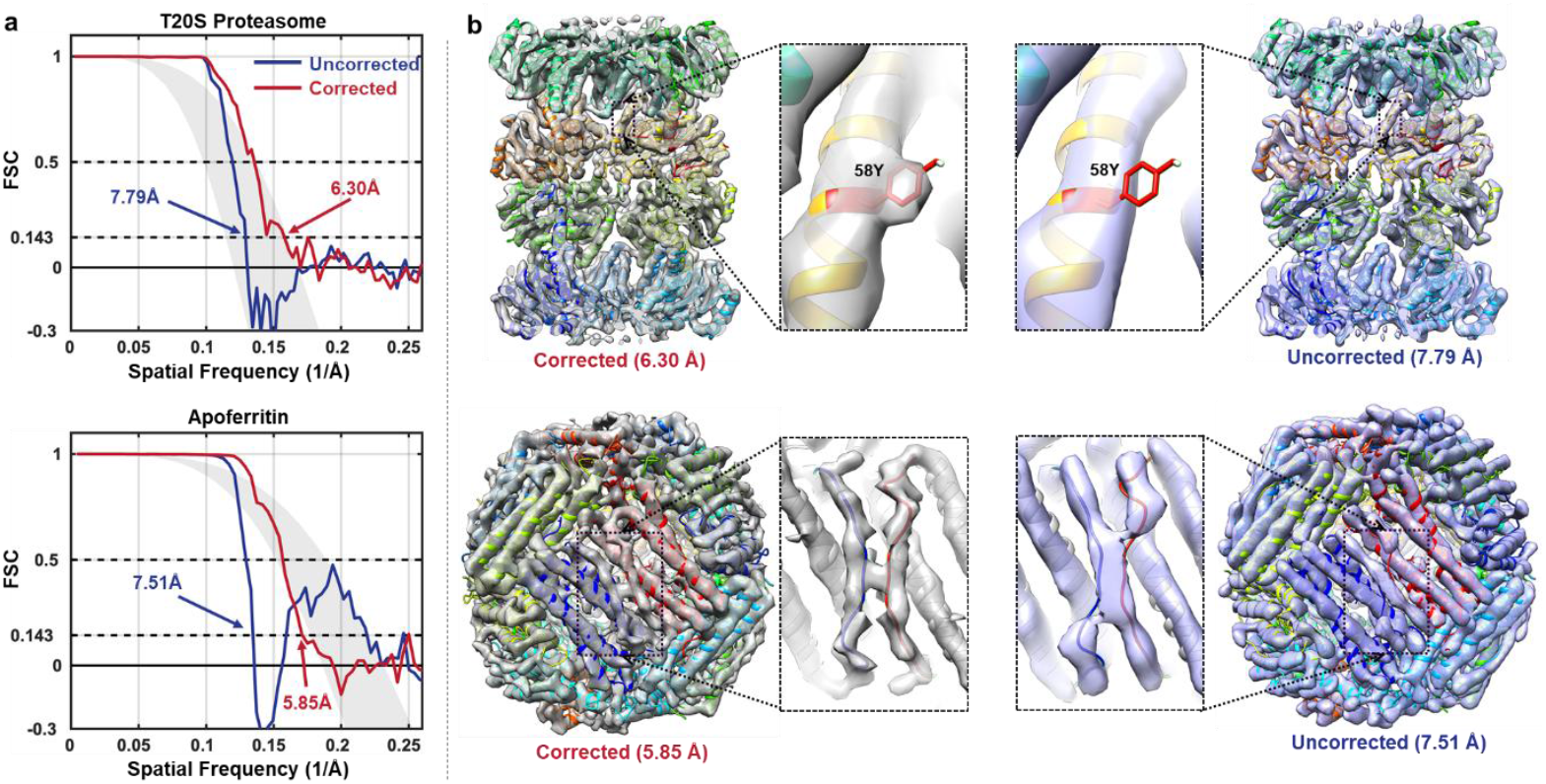
Comparisons of FSCs and density maps reconstructed by ptychographic SPA. The T20S proteasome (the upper row) and apoferritin (the lower row) datasets are used for the comparison. **a)** FSC curves of the 3D reconstructions from the corrected (red lines) and uncorrected (blue lines) micrographs. The gray bands indicate the range of varying cos(*χ*) in the defocus range of all particles. **b)** Density maps reconstructed from the corrected (left) and uncorrected (right) micrographs. Enlarged views of the local regions highlight a visible bulky side chain (residue 58Y) on the β-subunit of the T20 proteasome and two adjacent loops (residues between 78G and 94E) on the apoferritin.

The resolution improvements were subsequently verified using enhanced map features (**Figure 5b**). The corrected reconstruction of T20S proteasome revealed a bulky side chain (labeled with residue name in **Figure 5b**) that were absent in the uncorrected reconstructions. Two adjacent loops on the surface of apoferritin with a minimal distance of ∼5 Å (**Supplementary Figure 6a**) became distinctly separated after improving the overall resolution from 7.51 Å to 5.85 Å (**Figure 5b**). Although the published reconstruction from the same apoferritin dataset using CryoSPARC led to a 5.8 Å resolution with optimized processing, the two loops remained unresolved and resembled our uncorrected 7.51 Å resolution map (**Supplementary Figure 6b**). To eliminate any potential biases from software differences and processing optimization, we employed all particles selected by 2D classifications for THUNDER’s 3D reconstruction without additional particle selection while simultaneously processing the same particles with CryoSPARC and RELION using their respective optimal settings (**Supplementary Figure 6c and Supplementary Figure 7**). All the reconstructed results corroborated the findings obtained using THUNDER. Collectively, these comparisons demonstrate that the sampling mismatch constitutes a fundamental resolution limitation for ptychographic SPA.

## Discussion

In this study, we identified and investigated the impact of a sampling mismatch on ptychography, particularly its effects on ptychographic SPA. The sampling mismatch primarily arises from the mismatch between the pixel size of the reconstructed image patches at each scanning position and the scanning step size. This mismatch prevents the image patches from consistently aligning within overlapping regions. Owing to the inherent adaptive nature of ptychographic algorithms, this mismatch issue is not readily observable, but manifests as size deviations and a mismatch-induced modulation function (MIMF) that alters the information transfer in the final reconstructed micrographs. The primary consequence of MIMF is the introduction of zero-crossing oscillations in the Fourier domain of the potential function or micrograph extracted from the transmission function, which leads to phase reversals at certain spatial frequencies. When multiple micrographs are merged, as in SPA, phase reversals interfere destructively, leading to a loss of information and resolution. These effects were experimentally verified using two representative biological specimens. These findings demonstrate that eliminating sampling mismatches is crucial for achieving a high-resolution ptychographic SPA.

Sampling mismatch can arise in all applications of ptychography; however, we particularly emphasize its impact on the SPA of biological specimens. Biological imaging requires STEM to scan submicron or even larger areas to achieve sufficient imaging efficiency or accommodate the large size of biological structures. Such scanning ranges are far greater than those used in conventional materials-science experiments^1,2^, where the field of view is typically only tens of nanometers and calibrated using crystals under relatively high radiation doses. In contrast, the large scan areas required for biological samples make the accurate calibration of the step size difficult, especially given the lack of radiation-resistant and dimensionally precise standard samples. Ptychographic SPA also provides a method for correcting the scanning step size by matching the reconstructed density map of a known protein to its atomic model.

The phase reversal observed in the present work provides insight into how inaccurate sampling parameters affect the obtainable resolution, especially for ptychographic SPA. The first zero crossing of the real part of the MIMF provides an effective metric, as it identifies the highest spatial frequency unaffected by mismatch-induced signal confusion and thus approximates the upper bound of the achievable resolution. At a given defocus of 1.5 μm (**Figure 6a**), achieving the near-atomic resolution at 3.5 Å requires a mismatch less than 2% (*α*_s_*α*_f_ = 0.98), and the atomic resolution at 1.0 Å requires 0.2% (*α*_s_*α*_f_ = 0.998). Accordingly, if the pixel size of the CBED images can be calibrated accurately, the deviation in the scanning control or scanning step size must be controlled with similar precision. Meanwhile, we observed that a lower defocus may provide better tolerance for the sampling mismatch under a given target resolution (**Figure 6b**). However, a large defocus is often used in practice to improve the data collection efficiency. Therefore, balancing resolution and efficiency may be necessary in the presence of a sampling mismatch. These findings highlight the need for the precise calibration of sampling parameters and accurate control of scanning for high-resolution ptychographic SPA.

**Figure 6.**
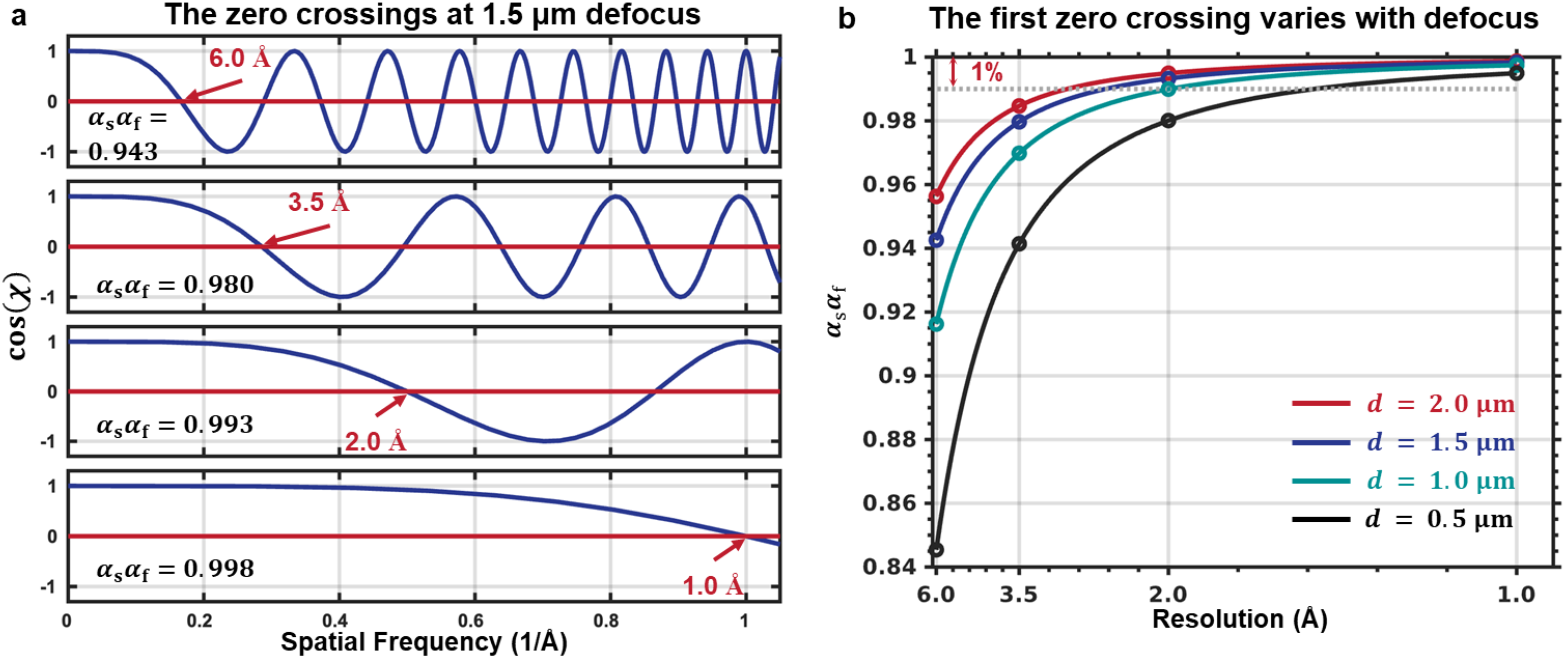
The zero crossings in the cos(*χ*). The real parts of the MIMF, cos(*χ*), presented as an information modulator for the micrographs, are compared under different sampling-mismatch factors and defocuses. **a**) cos(*χ*) under various sampling-mismatch factors at a fixed defocus of 1.5 μm. The positions of the first zero crossing are pointed with the corresponding resolution values. **b**) Relationship between the sampling-mismatch factors and the corresponding resolutions of the first zero crossing under various defocuses.

Additionally, the defocus measurement is affected by the presence of a sampling mismatch, which leads to two issues. First, the measured defocus may differ from actual physical values. Second, the defocus used in ptychography must be compatible with the sampling parameter deviations (**Eq. 2**). The variation in defocus with sampling parameter deviations revealed an intrinsic adaptive adjustment within the ptychographic calculation. Therefore, we need to re-estimate the defocus during testing with various deviations in the sampling parameters.

Although ptychography is theoretically capable of achieving extremely high resolutions, as demonstrated in materials science applications, its resolution on large biological specimens remains limited to the sub-nanometer scale. To the best of our knowledge, the resolution achieved in our current work is the highest one achievable using the ptychographic SPA. However, the factors preventing ptychographic SPA from reaching an even higher resolution remain unclear. A key step toward this goal is to clarify the underlying mechanism of information transfer. In this study, by treating ptychographic reconstruction as a virtual optical system, we derived the MIMF, which may offer novel insights into information transfer in computational imaging and guide future efforts to improve the resolution. Existing studies^6,7^ suggest that one major factor limiting the current resolution is beam-induced particle motion or, equivalently, inaccuracies in the scanning positions. Within the framework of this study, such mechanical jitter and sample drift can be viewed as another form of image-patch mismatch in reconstructing the overall transmission function, parallel to the sampling mismatch examined here. Therefore, the same conceptual framework may be extended to analyze scanning position inaccuracies in the future.

In this study, we employed two datasets to analyze the impact of sampling mismatches. The T20S proteasome dataset was collected using a first-generation Titan Krios microscope and an EMPAD detector. A potential factor preventing our T20S proteasome data from reaching the same resolution as the apoferritin data is the relatively long dead time of ∼0.8 milliseconds in the EMPAD detector. Given that the dwell time per scan point in our current setup is 1.8 milliseconds, this dead time means that out of the total accumulated dose of ∼63 e/Å^2^, nearly half the dose (2 8 e/Å^2^) was lost. T hus, this is a key factor resulting in a lower resolution of the T20S data than that of the apoferritin data.

In summary, the discovery and correction of sampling mismatches should play a significant role in further improving the resolution of ptychographic SPA and understanding information transfer in ptychographic computations.

## Methods

### Estimation of the modulation function from experimental transmission functions

To validate that the observed discrepancy between the calculated transmission functions 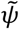 affected and *ψ* unaffected by the sampling mismatch stems from MIMF, we extracted the discrepancy as a function termed *H* from the experimental transmission functions. *H* should be a complex modulation function in the Fourier space, defined as 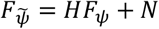, where *N* denotes additive noise. We assume that *H* is isotropic with respect to the angle. A least-squares problem was formulated within each Fourier ring at spatial frequency *u* with radial thickness Δ*u*, as

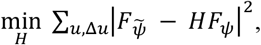

where ∑_*u*,Δ*u*_(·) denotes the summation over all points within the corresponding Fourier ring. Taking the derivative with respect to *H*^∗^ (using the Wirtinger derivatives) and setting it to zero yields

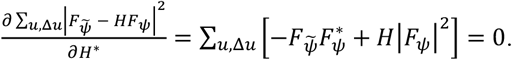

Rearranging yields

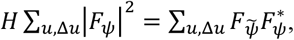

and solving for *H* gives the least-squares estimator

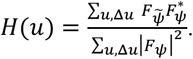

Then, *H*(*u*) was used to compare with the MIMF.

### FRC formula derived from MIMF

FRC was employed to analyze the phase reversals induced by sampling mismatches in the micrographs. In the following section, we establish the relationship between the FRC and the MIMF to interpret the observed phase reversals in the FRC, along with other additional features present in the FRC.

The transmission function obtained in ptychography can be modeled under weak-phase-object approximation as *ψ* = *e*^i*φ*^ ≈ 1 + i*φ*, where *φ* is the projection potential function. Taking the imaginary part of *ψ*, we can get the Fourier transform of the micrograph unaffected by the sampling mismatch as *F*_*φ*_.

Calculating the Fourier transform of *ψ*, we get *F*_*ψ*_ = *δ* + i*F*_*φ*_, where *δ* is the Dirac function. Substituting *F*_*ψ*_ into **Eq. 4** yields

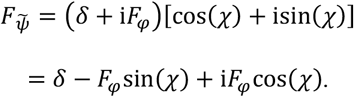

Since *F*_*φ*_ is the Fourier transform of the real-valued image and obeys conjugate symmetry, and cos(*χ*) and sin(*χ*) are even functions, *F*_*φ*_sin(*χ*) and *F*_*φ*_cos(*χ*) preserve the conjugate symmetry. Therefore, the inverse Fourier transforms of both *F*_*φ*_sin(*χ*) and *F*_*φ*_cos(*χ*) are real-valued images, corresponding to the real and imaginary parts of the sampling-mismatch-affected transmission function 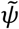. Consequently, we obtain 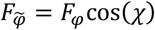, where 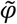 denotes the projection potential function affected by the sampling mismatch.

The FRC of **Eq. 3** can be written in the presence of noise as

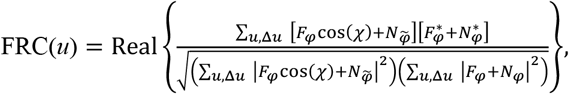

where 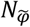 and *N*_*φ*_ are noises in the Fourier space for 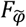 and *F*_*φ*_, respectively.

Assuming that the noises 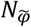 and *N*_*φ*_ are zero-mean and independent noise in a Fourier ring (uncorrelated with the signal), respectively, we can further simplify the FRC(*u*) above. The numerator becomes

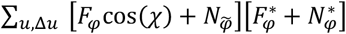

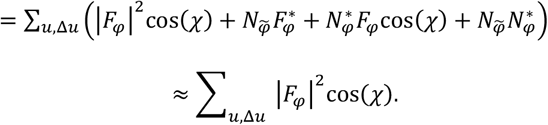

Similarly, the denominator becomes

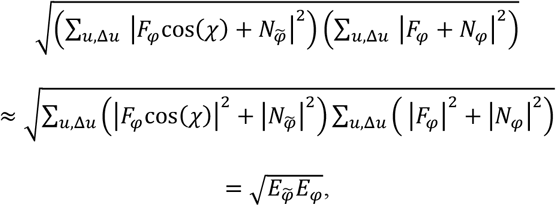

where 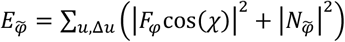 and *E*_*φ*_ = ∑_*u*,Δ*u*_ ( |*F*_*φ*_|^2^ + |*N*_*φ*_|^2^ ) represent the total power in the Fourier ring at spatial frequency *u*. Under the low-dose condition of the biological applications, the noise should be much stronger than the signal, and hence 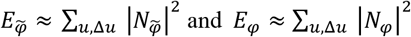 and *E*_*φ*_ ≈ ∑_*u*,Δ*u*_ |*N*_*φ*_|^2^ . The denominator represents the noise level in the Fourier rings.

Finally, we get the FRC formula from the MIMF

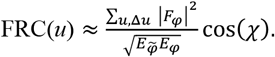

### Sample preparation and data acquisition of T20S proteasome

A drop of 4 μL of the purified T. Acidophilum 20S (T20S) proteasome at a concentration of ∼3 mg/mL was applied to glow-discharged Quantifoil holey carbon grids (Quantifoil Micro Tools GmbH) and plunge-frozen using Vitrobot Mark III (FEI Company).

The ptychographic dataset of the T20S proteasome was acquired using a 300 kV Titan Krios G1 (FEI Company) equipped with an Electron Microscope Pixel Array Detector G1^23^ (EMPAD G1, 128 × 128 pixels, Thermo Fisher Company) and a STEM system. A 20 μm C2 aperture was used, corresponding to a convergence semi-angle of ∼3.69 mrad. The probe was first brought into the in-focus condition on an amorphous carbon support film by optimizing the Ronchigram. Once the in-focus condition was established, the probe defocus was intentionally shifted by +1 µm for the ptychographic data collection over the areas of interest. Specifically, nine regions of interest were defined within each microhole of the ice-embedded sample, with each region covering a field of view of 305 nm (nominal value: 335 nm).

CBED images were recorded with ∼1.0-2.0 µm defocus and a nominal reciprocal-space pixel size of 0.005570 Å^-1^. A total of 388 micrographs were acquired in two sessions. In the first session, the scanning array was set to 300 × 300 points to cover a field of approximately 305 nm. In total, 222 micrographs were obtained. In the second session, the scanning array was set to 250 × 250 points to cover a field of approximately 305 nm. In total, 166 micrographs were obtained. Throughout these experiments, the electron dose was maintained at a consistent level, yielding a total accumulated dose of ∼63 e^−^/Å^2^. Due the dead time of EMPAD, nearly half of the total dose, ∼28 e^−^/Å^2^, was not recorded.

### Ptychographic micrograph reconstructions

Pre-processing of the T20S proteasome and apoferritin datasets^7^ was performed using the py4DSTEM software package^24^, including shift and rotation correction and defocus estimation by the tcBF-STEM method^16^.

Ptychographic reconstructions were performed using a batched stochastic gradient descent algorithm^17^ implemented in a custom MATLAB program (see Code Availability). Similar to the ePIE algorithm^9^, our code uses an iterative scheme that simultaneously updates the transmission and probe functions to minimize the discrepancy between the calculated and experimental CBED images. The iterative reconstruction was initialized with a unity-valued object function and an initial probe generated from the estimated defocus and probe-forming aperture. The probe-forming aperture was obtained by averaging all CBED images in the dataset. The algorithm was executed for over 30 iterations to achieve convergence.

Two sets of micrographs with corrected and uncorrected sampling parameters were generated for the T20S proteasome and apoferritin datasets using the methods described above.

### Single-particle analysis

Four sets of micrographs of the T20S proteasome and apoferritin datasets were processed using corrected pixel sizes (**Table 1**). The particles were picked using EPicker^25^ and then subjected to two rounds of screening through 2D classification using THUNDER^18^. The selected particles (**Table 1**) were recentered based on the shift from the image center of the corresponding 2D class averages and re-extracted from the corresponding micrographs. New particles from the four sets of micrographs were independently subjected to 3D reconstruction using THUNDER, cryoSPARC4.7.1^21^, and RELION5.0^22^. CTF correction in the software was turned off. The initial models were generated from atomic models of the T20S proteasome (PDB entry: 1PMA^19^) and apoferritin (PDB entry: 8RQB^7^) with the corresponding corrected pixel sizes (**Table 1**).

With THUNDER, all particles were used for 3D refinements starting at 60 Å resolution. Particle grading options were enabled during 3D reconstruction. The final reconstructions were then sharpened using an integrated postprocessing subroutine. The residue 58Y on β subunit of atomic model (PDB entry: 1PMA) of the T20S proteasome was manually fitted to the sharpened map of the corrected reconstruction using Coot^26^.

With cryoSPARC, the particles were further filtered by heterogeneous refinement for two classes with an initial model from *ab init* 3D model generation. The remaining particles were subjected to homogeneous refinement. The final reconstructions were processed using an integrated sharpening subroutine.

With RELION, all the particles were subjected to 3D auto-refinement starting at 40 Å resolution. The final reconstructions were sharpened using an integrated post-processing subroutine.

Chimera^27^ was used to prepare the figures.

### Pixel size calibration

To determine the pixel size of the 3D reconstruction, a reference map with the nominal pixel size was first generated from the corresponding atomic model using Chimera. The reconstructed and reference maps were aligned in the same orientation using the aligned 3D map utility in cryoSPARC. The EMDA software package^20^ was used to perform magnification refinement, yielding a scale factor that describes the relative magnification of the reconstructed map with respect to the reference. Finally, the corrected pixel size of the reconstructed map was obtained by dividing the nominal pixel size by this scale factor.

## Data availability

The datasets used and analyzed in the current study included the ptychographic dataset of apoferritin (EMPIAR-12236), the reconstructed map of apoferritin (EMD-19425), the atomic models of apoferritin (PDB:8RQB) and T20S Proteasome (PDB:1PMA). The T20S proteasome dataset acquired using an uncorrected cryo-electron microscope is available at EMPIAR-13137, and the maps obtained with the corrected and uncorrected sampling parameters are available at EMD-67291 and EMD-67292.

## Code availability

The custom MATLAB pipeline for ptychographic reconstruction is available at https://github.com/li-ty22/ptycho-workstation. The data-processing scripts utilized here are available at https://github.com/li-ty22/Sampling-Mismatch-and-Correction-for-Ptychographic-Single-Particle-Analysis.

## Acknowledgements

This work was supported by funds from the National Natural Science Foundation of China (32430056 for XL), National Key Research and Development Program (2024YFA1307302 for XL), Tsinghua-Peking Joint Center for Life Sciences, Beijing Frontier Research Center for Biological Structure, Shenzhen Medical Academy of Research and Translation (SMART). We acknowledge Ziying Zhang from Tsinghua University for guidance on sample preparation and single particle analysis; Jinying Ma and Qiaoyang Ma from Tsinghua University for their assistance and guidance in microscope operation; Fande Yu from Tsinghua University for their assistance with data acquisition; Zhao Wang and Mingtao Huang from Tsinghua University for technical support and helpful discussions. We acknowledge the Tsinghua University Branch of China National Center for Protein Sciences Beijing for providing facility support in computing and CryoEM instruments.

## Author contributions

X.L. and Y.S. initialized the project. T.L. and X.L. conducted the systematic investigation of sampling mismatch. Z.Y. prepared the T20S proteasome sample. S.L. designed the data-acquisition strategy. S.L., T.L., Z.Y. jointly carried out the data acquisition for the T20S proteasome dataset. T.L. designed the data-processing workflow, implemented the program and performed all ptychographic reconstructions. X.L. and T.L. conducted the single particle analysis. X.L. and T.L. wrote the manuscript. All the authors revised the manuscript.

## Supplementary Figures and Legends

**Supplementary Figure 1.**
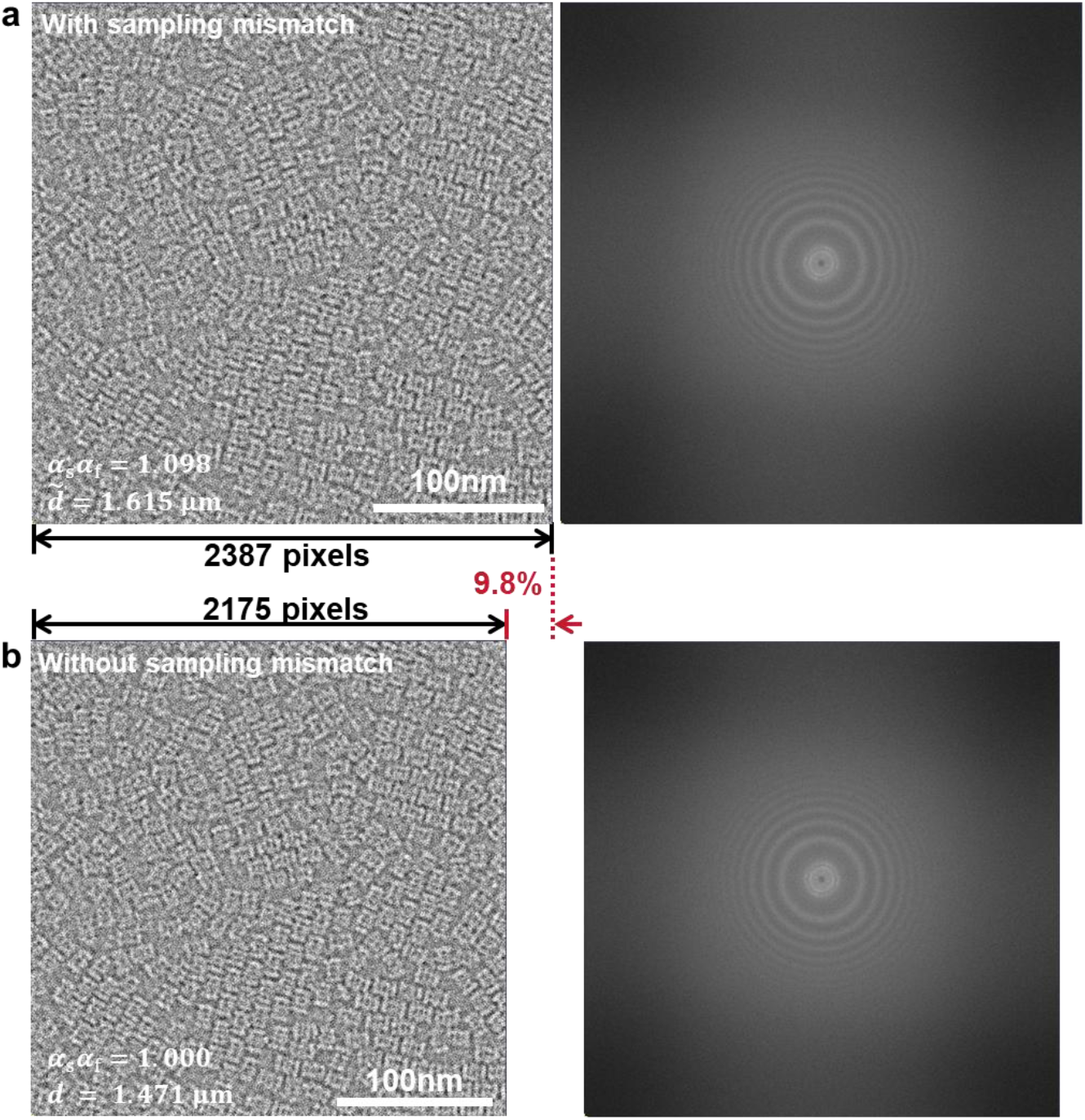
Typical micrographs in the T20S proteasome. A typical micrograph in the T20S proteasome dataset was shown. **a**) The micrograph reconstructed with sampling mismatch using the nominal scanning step size 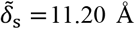 ( *α*_s_*α*_f_ = 1.098 ). **b**) The micrograph reconstructed without sampling mismatch using the corrected scanning step size *δ*_s_ = 10.20 Å ( *α*_s_*α*_f_ = 1.000 ). The power spectrums of the micrographs are shown on the right side. The sampling mismatch affects the final number of pixels in the micrograph.

**Supplementary Figure 2.**
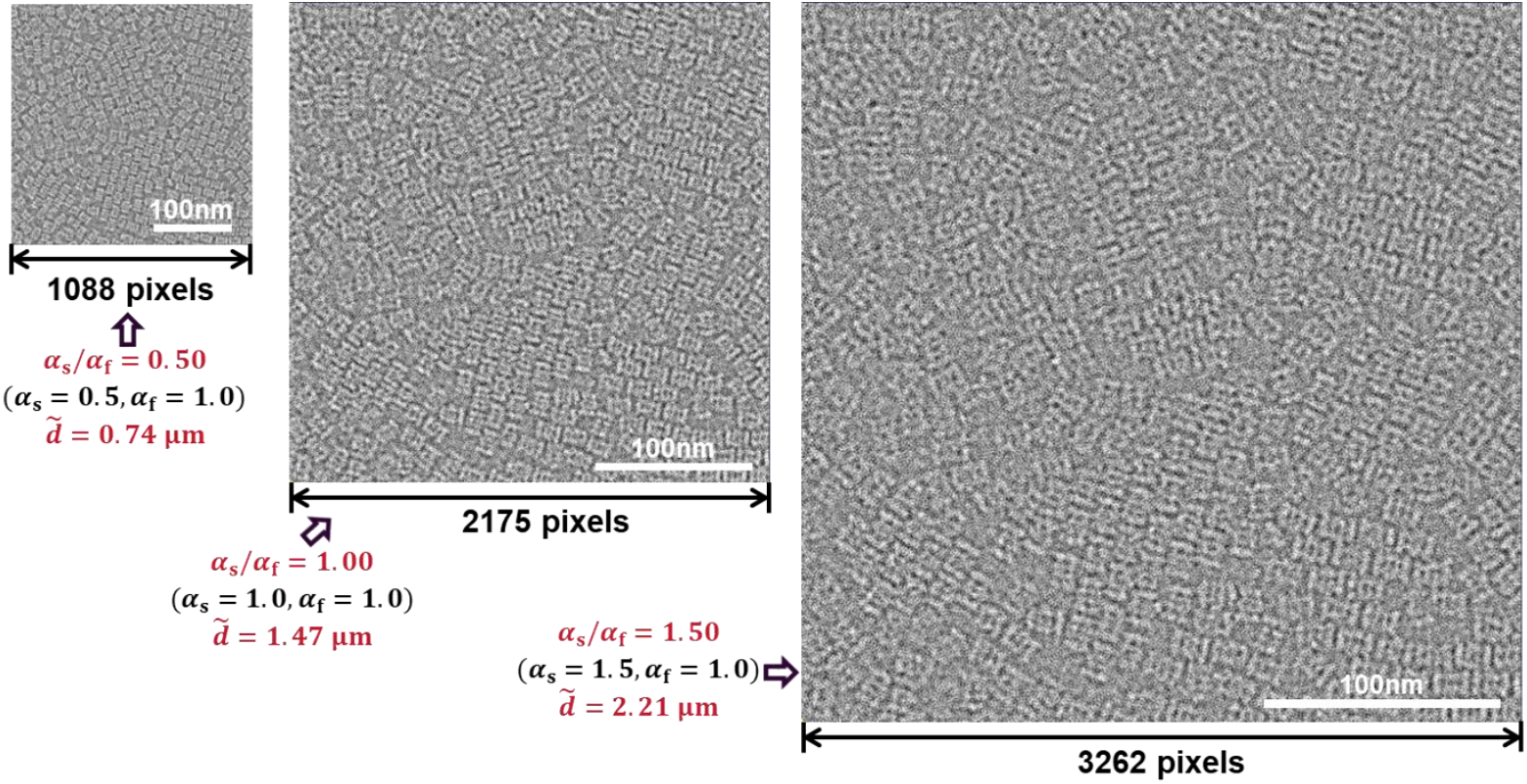
Size deviation induced by the sampling mismatch. A significant size deviation (the number of pixels shown below the micrograph) of the reconstructed micrographs can be observed under various sampling-mismatch factors. The corresponding defocus values (red) were measured. The set of CBED images corresponding to micrograph 2 in **Figure 2b** was used for the demonstration.

**Supplementary Figure 3.**
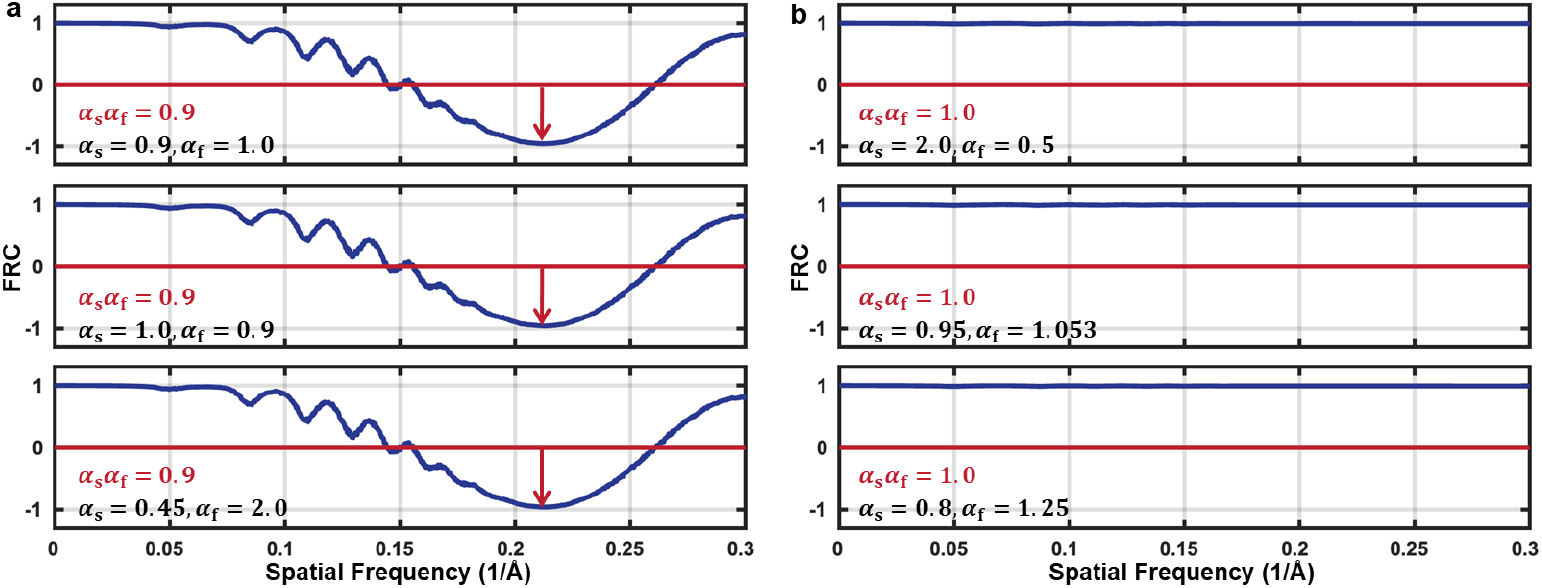
Unchanged micrographs under the same sampling-mismatch factor. The micrographs generated with two sampling-mismatch factors, **a**) 0.9 and **b**)1.0, but various combinations of *α*_s_ and *α*_f_, were used to calculated FRC curves. All micrographs were reconstructed from the same set of CBED images acquired at a defocus of 1.009 µm in the T20S proteasome dataset. The nearly identical FRC curves indicate that the signal differences among micrographs depend solely on *α*_s_*α*_f_. When *α*_s_*α*_f_ = 1, FRC becomes a constant 1, indicating no sampling mismatch.

**Supplementary Figure 4.**
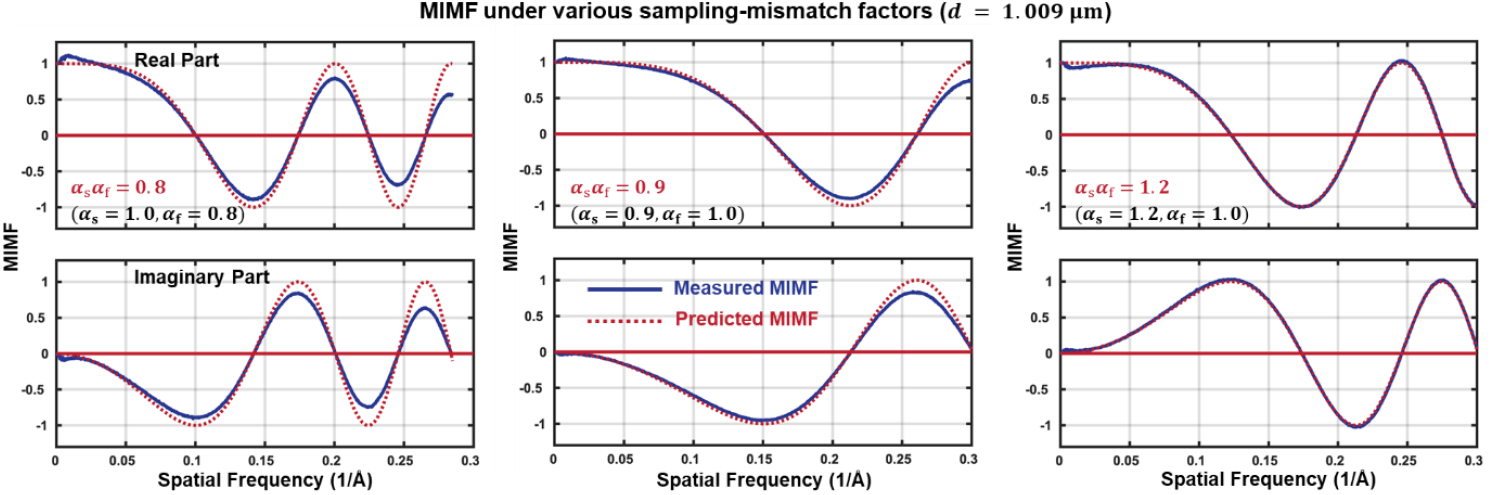
Examples of MIMF. Measured (solid blue) and predicted (red dashed) MIMF curves are shown for three sampling-mismatch conditions under the physical defocus *d* = 1.009 μm . Real and imaginary components of MIMF are displayed in the upper and lower panels, respectively.

**Supplementary Figure 5.**
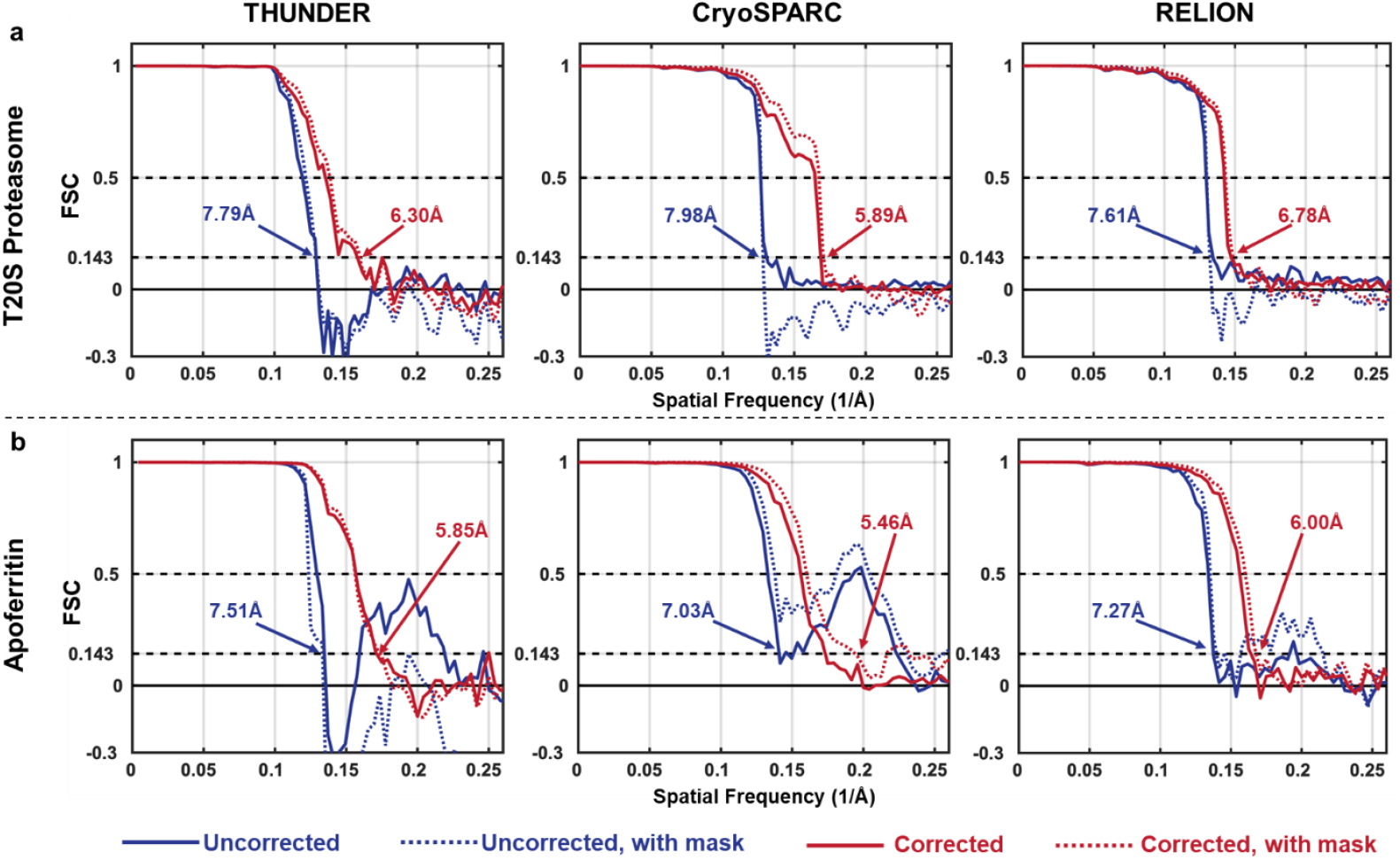
FSC curves from different processing software. FSC curves of two datasets, (**a**) T20S Proteasome and (**b**) apoferritin, processed by THUNDER, cryoSPARC, and RELION are plotted. FSC curves correspond to the dataset with uncorrected (blue) and corrected (red) sampling parameters are shown. For each reconstruction, FSC curves measured without (solid lines) and with (dashed lines) a mask are also shown. The resolutions were measured with a tight mask under the FSC criterion of 0.143 and listed in **Table 1**.

**Supplementary Figure 6.**
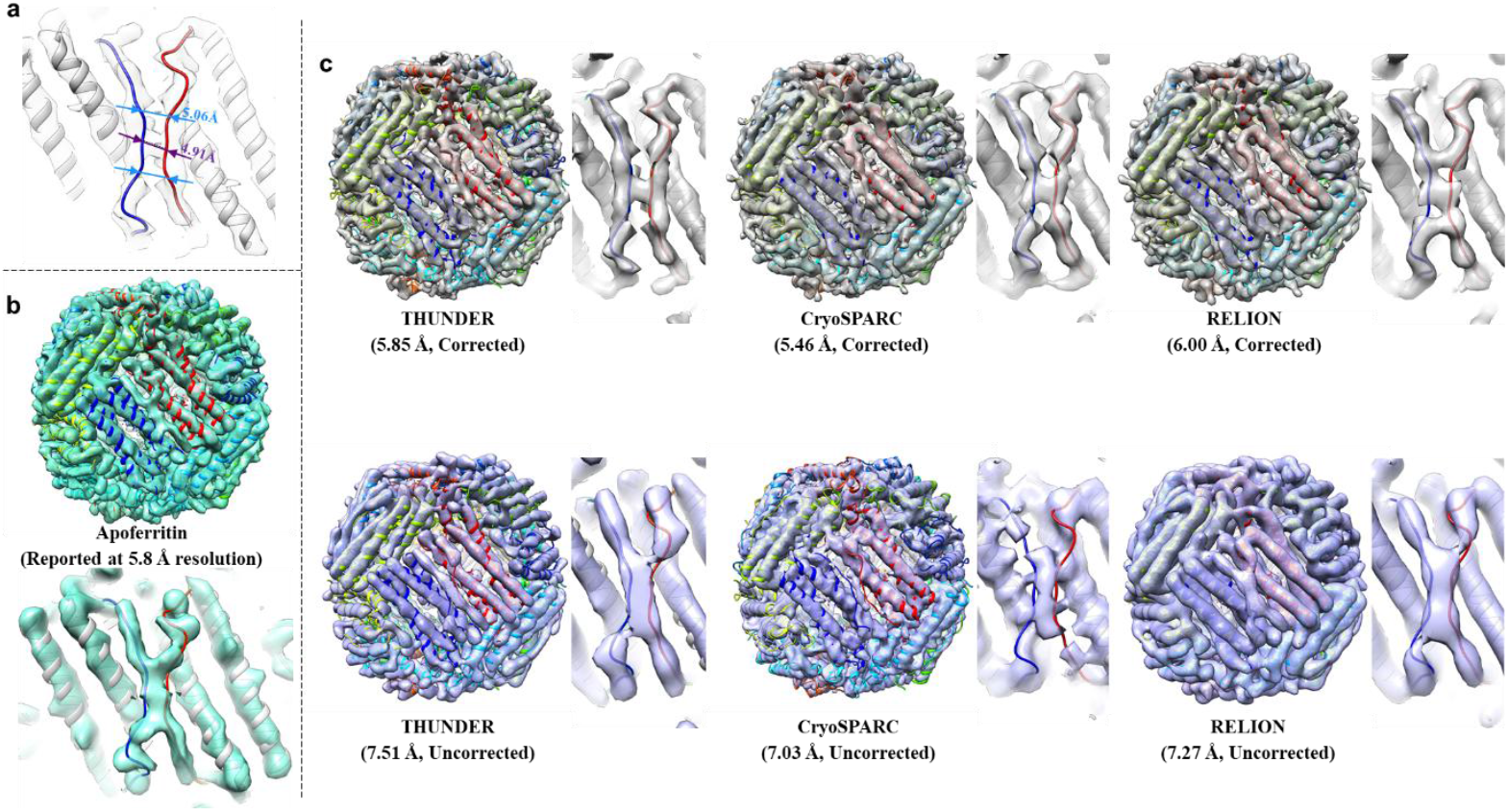
All density maps reconstructed from the apoferritin dataset. **a**) Two loops on the surface of apoferritin were used as a resolution indicator, which have a minimal distance of ∼5 Å. **b**) The originally published reconstruction (EMD-19425) of the apoferritin dataset (EMPIAR-12236). The densities of the two loops are not well separated in the middle region. **c**) Density maps of the whole apoferritin and two surface loops reconstructed with the corrected (upper row) and uncorrected (lower row) sampling parameters using THUNDER, CryoSPARC, and RELION. The atomic model (PDB: 8RQB) was shown with the maps. The resolutions were shown below the maps. The corresponding FSC curves and measured resolution values are summarized in **Table 1** and **Supplementary Figure 5**.

**Supplementary Figure 7.**
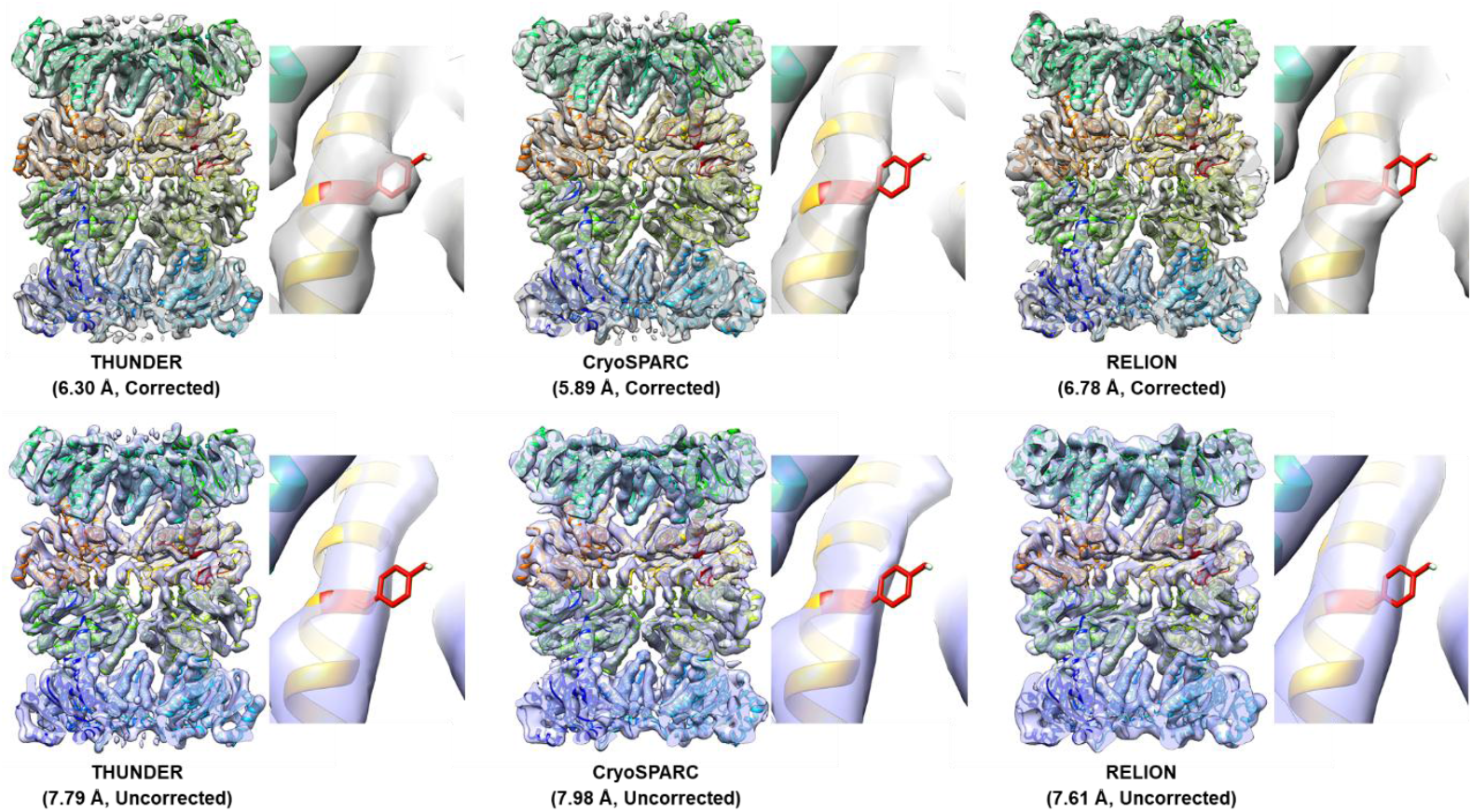
All density maps reconstructed from the T20S Proteasome dataset. Density maps of the whole T20S proteasome and a bulky size chain on the β subunit are shown, which are reconstructed with the corrected (upper row) and uncorrected (lower row) sampling parameters using THUNDER, CryoSPARC, and RELION, respectively. The fitted model based on the published one (PDB: 1PMA) is shown with the maps. The resolutions were shown below the maps. The corresponding FSC curves and measured resolution values are summarized in **Table 1** and **Supplementary Figure 5**.

